# High Coverage Highly Accurate Long-Read Sequencing of a Mouse Neuronal Cell Line Using the PacBio Revio Sequencer

**DOI:** 10.1101/2023.06.06.543940

**Authors:** Juana G. Manuel, Hillary B. Heins, Sandra Crocker, Julie A. Neidich, Lisa Sadzewicz, Luke Tallon, Tychele N. Turner

**Affiliations:** Department of Genetics, Washington University School of Medicine, St. Louis, MO 63110, USA; Department of Pathology and Immunology, Washington University School of Medicine, St. Louis, MO 63110, USA; Department of Pediatrics, Washington University School of Medicine, St. Louis, MO 63110, USA; Institute for Genome Sciences, University of Maryland School of Medicine, Baltimore, MD 21201, USA

**Author notes:** Correspondence to Tychele N. Turner, Ph.D., Washington University School of Medicine, Department of Genetics, 4523 Clayton Avenue, Campus Box 8232, St. Louis, MO 63110. co-first authors (Juana G. Manuel, Hillary B. Heins).

**Keywords:** Pacific Biosciences, PacBio, Revio, HiFi, long-read whole-genome sequencing, long-read WGS, neuronal, cell line

## Abstract

Recently, Pacific Biosciences released a new highly accurate long-read sequencer called the Revio System that is projected to generate 30× HiFi whole-genome sequencing for the human genome within one sequencing SMRT Cell. Mouse and human genomes are similar in size. In this study, we sought to test this new sequencer by characterizing the genome and epigenome of the mouse neuronal cell line Neuro-2a. We generated long-read HiFi whole-genome sequencing on three Revio SMRT Cells, achieving a total coverage of 98×, with 30×, 32×, and 36× coverage respectively for each of the three Revio SMRT Cells. We performed several tests on these data including single-nucleotide variant and small insertion detection using GPU-accelerated DeepVariant, structural variant detection with pbsv, methylation detection with pb-CpG-tools, and generating *de novo* assemblies with the HiCanu and hifiasm assemblers. Overall, we find consistency across SMRT Cells in coverage, detection of variation, methylation, and *de novo* assemblies for each of the three SMRT Cells.

## BACKGROUND

Highly accurate long-read whole-genome sequencing (WGS) (1) has been available for several years, but until recently has been comparatively expensive for the generation of 30× human genome coverage relative to short-read sequencing. Within the last year, PacBio HiFi sequencing has expanded to a new sequencer (the Pacific Biosciences Revio System) that allows for the generation of a 30× human genome in one Revio SMRT Cell with a significant reduction in cost. The human genome is estimated to be 3.055 billion-base pairs (2) and the mouse genome is estimated to be 2.731 billion base pairs (https://www.ncbi.nlm.nih.gov/assembly/GCF_000001635.20/). We sought to test the sequencing capabilities on Revio and chose the mouse neuronal cell line Neuro-2a for this research assessment. Neuro-2a is a cell line that has been utilized in >1000 papers (based on a PubMed search on May 22, 2023) and in particular, has also been used in the ENCODE project (3) (https://www.encodeproject.org/search/?type=Biosample&searchTerm=neuro-2a) for studies of regulatory element function (4). Key analyses of long-read WGS include reference-based and assembly-based strategies. More recently, 5mC methylation detection has also become possible with long-read WGS (5). Herein, we describe our analyses of highly accurate long-read WGS of the Neuro-2a cell line.

## MATERIALS AND METHODS

### Cell Line

The Neuro-2a Mouse Neuronal Cell Line (ATCC CCL-131, lot # 70041794) at Passage 184 was purchased from ATCC and grown under standard conditions (EMEM medium [ATCC, catalog # 30-2003] + 10% FBS [Gibco 26140-079], and Pen-Strep [Gibco 15140122] in an incubator set at 37°C with 5% CO2).

### Karyotype

Neuro-2a cells were initially grown in a 100 mm plate. Cells were then transferred to a T25 flask (TPP 90026) and incubated at 37°C with 5% CO2. These adherent cells were allowed to reach 60 – 70% confluency. The Neuro-2a cells were sent to the Washington University in St. Louis Cytogenetics and Molecular Pathology Core Laboratory. The Neuro-2a cells were treated with a hypotonic solution, fixed, and then stained using the Giemsa (GTG) banding method. 20 metaphases were examined using a microscope and the CytoVision imaging software.

### PacBio WGS sample preparation and HiFi sequencing

High Molecular Weight (HMW) DNA samples were generated using Quick-DNA HMW MagBead Kit (Zymo Research D6060). Each sample was prepared following the provided protocol with certain changes. Approximately three million cells were used per sample, and 6-8 samples were combined after collection to acquire enough HMW DNA for sequencing. Liquid sample and Biofluid & Solid Tissue Buffer were changed to 300 μL each. Quick-DNA MagBinding Buffer was changed to 600μL, and MagBinding Beads were changed to 100 μL. When pipette mixing, we mixed with regular pipette tips 3 – 4 times to break up clumps and then used wide bore pipette tips to reduce shearing (Labcon 1199-965-008-9) to continue mixing. DNA Elution Buffer was changed to 150 μL per sample. Samples were combined for quantitation using Qubit dsDNA Broad Range Assay (Invitrogen Q32853). Sample was stored at 4°C to avoid freeze-thaw. The HMW DNA sample was sequenced on a PacBio Revio System sequencer at Maryland Genomics at the Institute for Genome Sciences, University of Maryland School of Medicine on three Revio SMRT Cells.

Post-sequencing, the data were analyzed using two main approaches: *de novo* assembly-based and reference-based workflows. For *de novo* assembly, two different assemblers were used, including HiCanu (6) version 2.2 and Hifiasm (7) version 0.16.1-r375. For the reference-based workflow, the reads were mapped to the mm10 genome (http://hgdownload.cse.ucsc.edu/goldenpath/mm10/bigZips/mm10.fa.gz) using pbmm2 (8) version 1.10.0 align, followed by merging of the data from three Revio SMRT Cells with SAMtools version 1.6 merge, sorting of the reads with SAMtools sort, and indexing of the bam with SAMtools index.

SNV/indel calls were generated using DeepVariant (9) implemented in NVIDIA Parabricks version 4.1.0-1. Structural variants were called using pbsv version 2.9.0. Post-calling SVs were merged using SURVIVOR (10) version 1.0.7 merge with these specific arguments to consider a max distance between breakpoints of 10 bp, take the type of SV into account, take the strand of the SV into account, and to examine SVs at least 50 bp in size. The SVs were then compared using SURVIVOR genComp. 5mC status was detected using pb-CpG-tools (11) aligned_bam_to_cpg_scores version 2.3.1 with the pileup_calling_model.v1.tflite model file and minimum mapq of 30, and minimum coverage of 10.

## RESULTS

### Karyotype reveals a complex aneuploid genome

Since the Neuro-2a genome is known to be an immortalized cell line and due to a previous genomic workup of this cell line (12), we hypothesized it may be aneuploid. The results of the karyotype analysis (**Figure 1**) confirmed this idea with a chromosome complement of 79∼97<4n>,X/Yx0,-2,-4,-3,-4,-5,-6,-7,-8,-8×2,-9,-10,-11,- 13×2,+14,+15×3,+16,+16×2,-18,+19×2,+9-23mar,1-5dmin[cp10]. This cell line was considered to be near tetraploid with 4n=80 and the karyotype features gains and loss of multiple chromosomes as well as no identifiable sex chromosomes. 23 marker chromosomes (including metacentric and submetacentric markers) and up to 5 double minutes (dmin) were observed. A marker chromosome is a structurally rearranged chromosome whose origin cannot be definitively defined. Double minutes (dmin) are acentric fragments that manifest from gene amplification which gives the cells selective advantages for growth and survival. They frequently harbor amplified oncogenes and genes involved in drug resistance.

**Figure 1.**
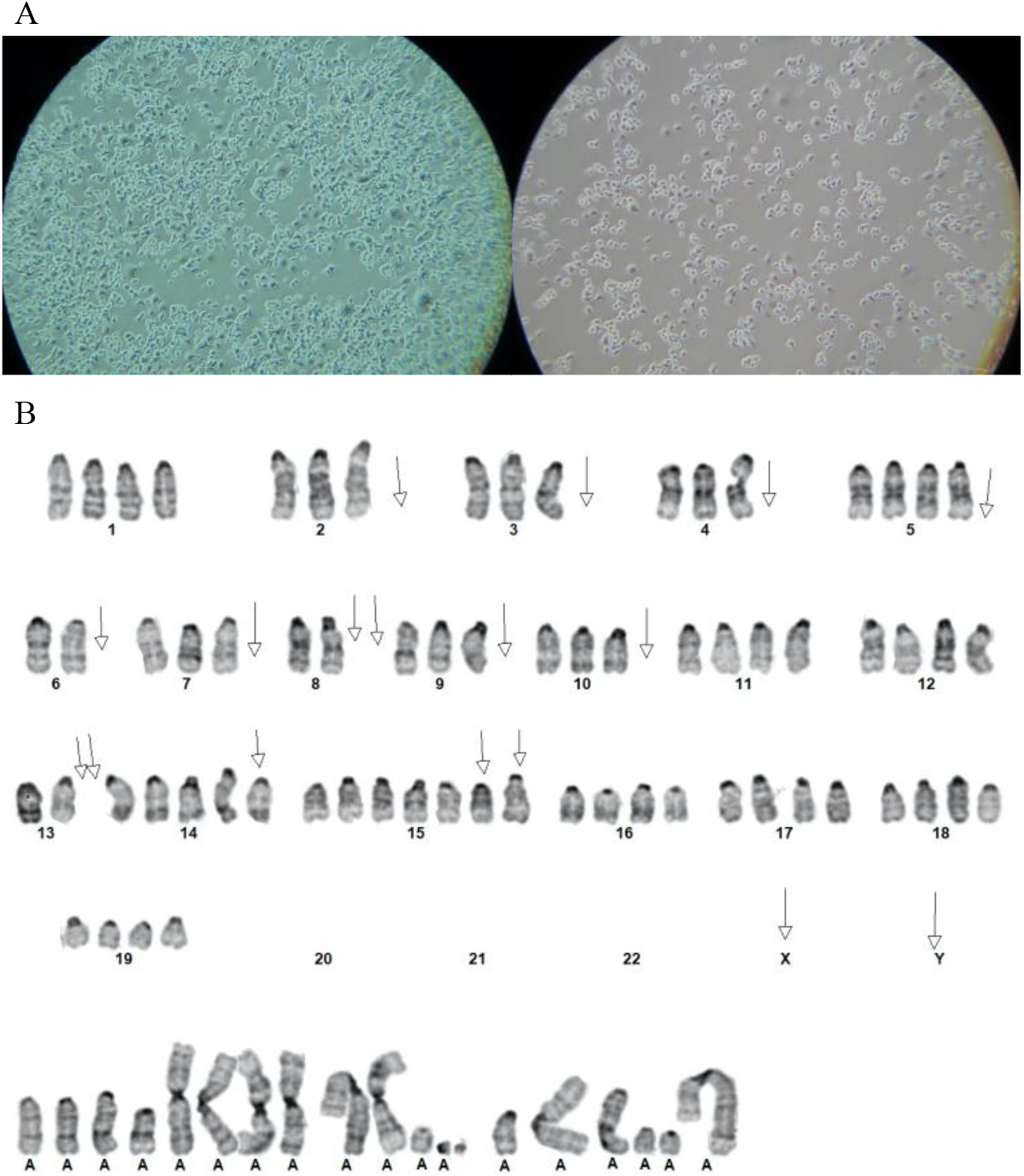
Karyotype of the Neuro-2a Mouse Hippocampal Neuronal Cell Line. (A) First picture is of Neuro-2a the three days after resurrecting and plating the cell line. The second picture is of Neuro-2a in a T25 flask on the day when we sent the cells for karyotyping. Pictures are taken at 10**×**. (B) The Karyotyping of Neuro-2a was done by the GTW banding method. Metaphase cells had the characteristic of chromosomally abnormal near tetraploid composite karyotype. During karyotyping, there is loss of both sex chromosomes, so since the known sex chromosomes had not been provided, the sex chromosomes could not be determined.

### Revio SMRT Cells are consistent in sequencing characteristics

Three PacBio Revio SMRT Cells were sequenced for the Neuro-2a cell line (**Table 1**). Each SMRT Cell collected kinetics data so that methylation could be detected in the downstream analyses. All three of the SMRT Cells performed within expectation generating 30×, 32×, and 36× genome coverage, respectively. The mean HiFi read length for the SMRT cells was 14,069 bp, 14,103 bp, and 14,066 bp, respectively. This is a bit shorter than the optimal HiFi read length of 20,000 bp. Notably, for computational considerations the file size of each SMRT Cell HiFi bam was 352 GB, 377 GB, and 421 GB, respectively. After mapping to the mm10 genome, the file sizes were 429 GB, 460 GB, and 516 GB, respectively. Combining all three SMRT Cells the genome coverage was 98× and the resulting file size was 1,405 GB.

**Table 1.**
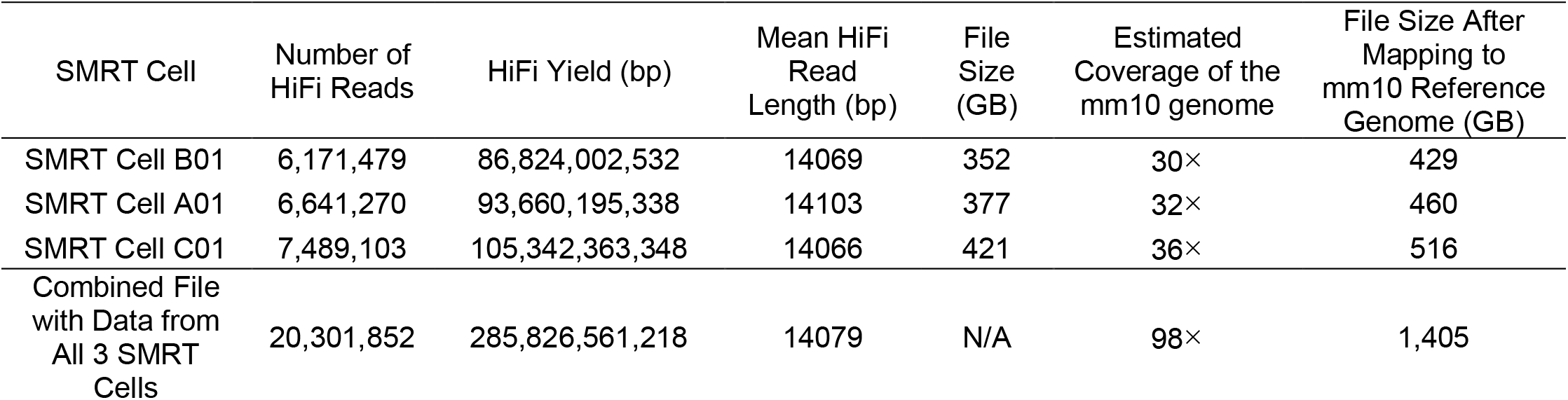
Sequencing metrics of the three PacBio Revio SMRT Cells

*SNV/indel calling, SV calling, and methylation detection are consistent across Revio SMRT Cells* Around the time we sequenced the Neuro-2a genome, NVIDIA also released a new version of their Parabricks software (13) for accelerating genomic analyses. This version (4.1.0-1) included rapid calling of SNVs/indels genome-wide from highly accurate long-read sequencing data. We previously have shown the advantage of GPU acceleration on short-read whole-genome sequencing using Parabricks (14) and sought to test the new version on our Revio HiFi data. For the analysis, we utilized a custom-built Lambda AI Vector Workstation. The workstation contains an AMD Threadripper Pro 5995WX Processer with 64 cores, and 2 NVIDIA RTX 6000 Ada GPUs. We ran Parabricks DeepVariant on each of the three SMRT Cells mapped to the mm10 genome independently and saw run times of 1422 seconds, 1542 seconds, and 1740 seconds, respectively (**Table 2**). Testing was also done on the full dataset at 98× coverage and the analysis ran in 4,585 seconds. This showed that for a 30× genome SNV/indel calling can be performed in <30 minutes. Furthermore, we performed concordance checks between each of the Revio SMRT Cells and the results from the full 98× coverage genome. The concordance, considering both SNVs and INDELs, was 97.9%, 98.0%, and 98.1%, respectively.

**Table 2.** SNV/INDEL calling in the three PacBio Revio SMRT Cells

| SMRT Cell | DeepVariant RunTime* | Concordance of SMRT Cell with Full Dataset** (%) | Number of SNVs | Number of INDELs |
| --- | --- | --- | --- | --- |
| SMRT Cell B01 | 1,422 | 97.9 | 7,215,187 | 2,754,428 |
| SMRT Cell A01 | 1,542 | 98.0 | 7,228,880 | 2,741,760 |
| SMRT Cell C01 | 1,740 | 98.1 | 7,255,687 | 2,668,512 |
| Combined File with Data from All 3 SMRT Cells | 4,585 | 100 | 7,441,882 | 2,500,894 |

\* Run on a Lambda AI Vector Workstation with an AMD Threadripper Pro 5995WX Processor with 64 cores, and 2 NVIDIA RTX 6000 Ada GPUs (seconds)
\*\* includes SNVs and INDELs

We compared SVs detected in the full dataset and in each of the Revio SMRT Cells (**Table 3**) and found high concordance with each pairwise comparison containing at least 88% of the SV calls. Finally, we compared the methylation values genome-wide in pairwise comparisons (**Table 4**) using a correlation test and found correlation for each pairwise comparison of at least 0.94 for each of the comparisons.

**Table 3.** PBSV concordance comparisons for the three PacBio Revio SMRT Cells

|  | Combined<br>File with Data<br>from All 3<br>SMRT Cells | SMRT Cell B01 | SMRT Cell A01 | SMRT Cell C01 |
| --- | --- | --- | --- | --- |
| Combined File with<br>Data from All 3<br>SMRT Cells | 1 | 0.892 | 0.887 | 0.897 |
| SMRT Cell B01 | 0.911 | 1 | 0.904 | 0.907 |
| SMRT Cell A01 | 0.908 | 0.905 | 1 | 0.907 |
| SMRT Cell C01 | 0.914 | 0.904 | 0.903 | 1 |

**Table 4.** Correlation of methylation values across three PacBio Revio SMRT Cells

|  | Combined<br>File with Data<br>from All 3<br>SMRT Cells | SMRT Cell B01 | SMRT Cell A01 | SMRT Cell C01 |
| --- | --- | --- | --- | --- |
| Combined File with<br>Data from All 3<br>SMRT Cells | 1 | 0.98 | 0.98 | 0.98 |
| SMRT Cell B01 | 0.98 | 1 | 0.95 | 0.95 |
| SMRT Cell A01 | 0.98 | 0.95 | 1 | 0.94 |
| SMRT Cell C01 | 0.98 | 0.95 | 0.94 | 1 |

### de novo assembly of the Neuro-2a genome

Two *de novo* assemblers (HiCanu (15) and hifiasm (7)) were utilized to test the assembly of the Neuro-2a genome. As noted, the karyotype is quite complex. For this reason, generating a *de novo* assembly for this cell line can present a challenge. The HiCanu assembler generated consistent assemblies for all three Revio SMRT Cells and a very similar size assembly even at 98× (**Table 5**). All assemblies were ∼2.8 Gbp in size with NG50 contig values between 53 and 64 and overall number of contigs between 1607 and 1877. The NG50 value ranged from 12,743,571 bp to 17,334,001 bp in the assemblies.

**Table 5.**
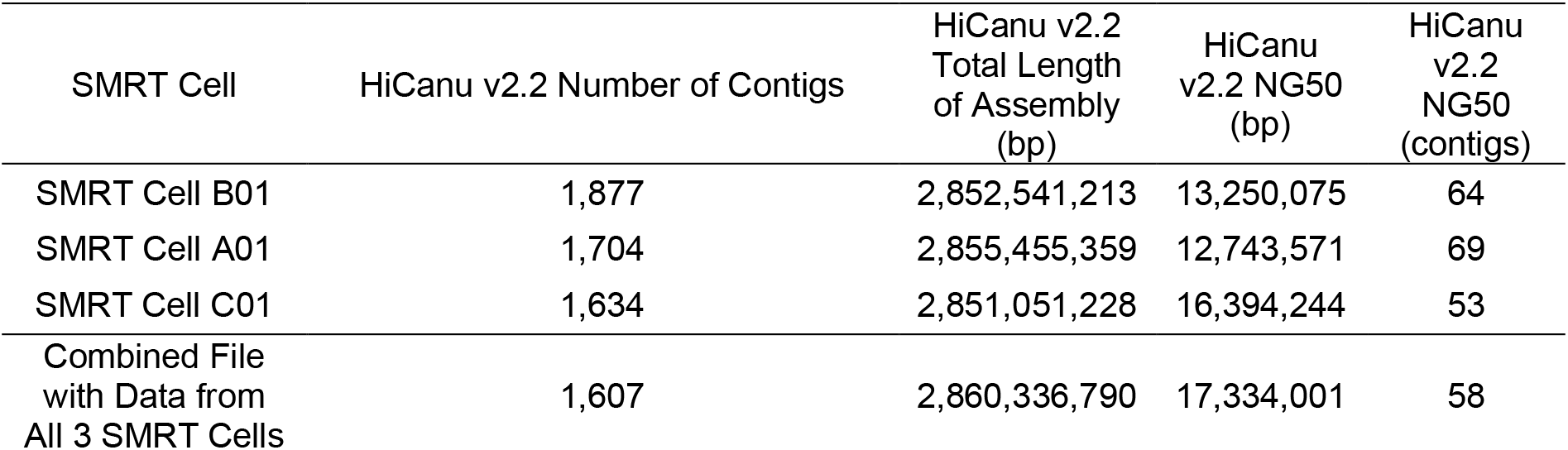
HiCanu assembly statistics for the three PacBio Revio SMRT Cells.

| SMRT Cell | HiCanu v2.2 Number of Contigs | HiCanu v2.2 Total Length of Assembly (bp) | HiCanu v2.2 NG50 (bp) | HiCanu v2.2 NG50 (contigs) |
| --- | --- | --- | --- | --- |
| SMRT Cell B01 | 1,877 | 2,852,541,213 | 13,250,075 | 64 |
| SMRT Cell A01 | 1,704 | 2,855,455,359 | 12,743,571 | 69 |
| SMRT Cell C01 | 1,634 | 2,851,051,228 | 16,394,244 | 53 |
| Combined File with Data from All 3 SMRT Cells | 1,607 | 2,860,336,790 | 17,334,001 | 58 |

The hifiasm assembler generated high-quality assemblies for all three Revio SMRT Cells (**Table 6**). We report statistics for both the hap1 and hap2 assemblies but since this genome is aneuploid we focus on characteristics of the primary assembly (.asm.bp.**p**_ctg). The assemblies ranged in size from ∼2.8 Gbp to ∼3.0 Gbp with NG 50 values between 33 and 44 and number of contigs ranging from 1,358 to 2,066. The NG50 value ranged from 21,996,525 bp to 26,122,379 bp in the assemblies.

**Table 6.** hifiasm assembly statistics for the three PacBio Revio SMRT Cells

| SMRT cell | hifiasm .asm.bp.ha p1.p_ctg.g fa.ctg NG50 (bp) | hifiasm .asm.bp.ha p1.p_ctg.g fa.ctg NG50 (contigs) | hifiasm .asm.bp.ha p2.p_ctg.gf a.ctg NG50 (bp) | hifiasm .asm.bp.ha p2.p_ctg.gf a.ctg NG50 (contigs) | hifiasm .asm.bp.p _ctg.gfa.ct g total contigs | hifiasm .asm.bp.p _ctg.gfa.ctg NG50 (bp) | hifiasm .asm.bp.p _ctg.gfa.ctg NG50 (contigs) |
| --- | --- | --- | --- | --- | --- | --- | --- |
| SMRT Cell B01 | 12,379,411 | 60 | 11,459,661 | 65 | 1,558 | 21,996,525 | 44 |
| SMRT Cell A01 | 11,538,520 | 63 | 11,562,341 | 66 | 1,358 | 22,705,572 | 38 |
| SMRT Cell C01 | 15,456,526 | 49 | 12,528,891 | 59 | 1,657 | 22,944,273 | 34 |
| Combined File with Data from All 3 SMRT Cells | 17,547,755 | 45 | 16,891,061 | 53 | 2,066 | 26,122,379 | 33 |

## DISCUSSION

Highly accurate long-read WGS on the PacBio Revio System is consistent and can generate 30× genome coverage on one SMRT Cell. We utilized this technology on a neuronal cell line to learn more about the characteristics of Revio HiFi sequencing data and to generate a genome and epigenome of the Neuro-2a cell line for research into neurodevelopmental and neurodegenerative disorders. Our analyses of these data showed consistency in small genomic variants, structural variants, methylation, and *de novo* assembly characteristics. For the *de novo* assemblies we did not observe a significant improvement in assembly quality using data from all three Revio SMRT Cells together. However, this is likely due to the complexity of the genome as evidenced by the complex karyotype (**Figure 1**).

## AVAILABILITY AND REQUIREMENTS

The data described in this preprint can be found at Globus link https://app.globus.org/file-manager?origin_id=4865823e-01af-11ee-a924-63e0d97254cd&path=%2F. Since the NCBI SRA does not have Revio as an option for a sequencing type, we have not yet uploaded the data there. We will upload the data there as soon as that is possible as part of BioProject PRJNA938057.

## ACKNOWLEDMENTS

Funding: This work was supported by a grant from the National Institutes of Health (R01MH126933). Author Contributions: T.N.T designed the study. J.M. and H.H. performed the cell culture work. S.C. and J.A.N. performed the karyotype. L.S. and L.T. performed the sequencing. T.N.T. analyzed the data. Data and materials availability: The data described in this preprint can be found at Globus link https://app.globus.org/file-manager?origin_id=4865823e-01af-11ee-a924-63e0d97254cd&path=%2F.

